# Uncovering the diversity of heterotrophic culturable bacteria within the gut of bay scallop *Argopecten purpuratus* (Lamarck, 1819) from two geographically distant natural banks

**DOI:** 10.1101/2023.12.04.569930

**Authors:** Wilbert Serrano, Ulrike I. Tarazona-Janampa, Raul M. Olaechea, Sonia Valle

**Author notes:** Universidad Científica del Sur, Antigua Panamericana Sur, Km.19, Lima-42. Author for correspondence: Wilbert Serrano.

## Abstract

We study the bacterial diversity within the gut of the peruvian bay scallop *Argopecten purpuratus* (Lamarck, 1819) (Mollusca: Pectinidae) collected from two shellfish growing areas in Independence Bay (IB) and Sechura Bay (SB) in Peru. This work aimed to identify isolates of culturable gut-associated microbiota of *A. purpuratus* through repetitive sequence-based PCR (rep-PCR) and 16S rRNA gene sequences comparison. A total of 20 healthy adult scallops were collected from each location twice, in spring and autumn, 2015 and 2016, respectively.

Fecal pellets from the distal intestinal tract of each individual were extracted, and after pooling, samples were cultivated on general and selective media following standard culture conditions. In order to prevent duplicate sequencing of the ribosomal 16S rRNA gene sequences of the isolates, a pre-screened step by (rep-PCR) amplification utilizing BOXA1R primer was performed, then isolates showing different banding profiles were selected for further sequencing.

Taxonomic distribution based on the 16S rRNA gene comparison of selected strains placed isolates from IB and SB into 29 and 27 different genera, representing 50 and 42 species, respectively. The major proportions of bacterial species found in IB corresponded to the division Firmicutes (46%), whereas the Proteobacteria and Firmicutes were almost equally distributed in (SB) with 38 and 43% respectively. Remarkable, (IB) showed a high number (28%) and diversity of species within the division Actinobacteria compared to (SB); while the species *Salinimicrobium* sp. of the division Flavobacteria was isolated only in (SB).

Common species in both areas are mainly related to the genus *Bacillus,* while strains of species *Vibrio alginolyticus* and *V. parahaemoliticus* were common representatives of the Proteobacteria. The intestinal tract of *A. purpuratus* may represent an uninvestigated niche for analyzing the structure of new complex microbial communities. This study showed that the diversity of culturable microbiota within the gut of two populations of *A. purpuratus* is significantly different at each location.

## INTRODUCTION

The bay scallop *Argopecten purpuratus* (Lamarck, 1819) (Mollusca: Pectinidae) represents the second invertebrate species of high commercial value in Peru. Natural environments are the primary source of the seeds of *A. purpuratus*, which further are massively cultivated in artificial fattening systems in bottom cages or suspended lanterns. The natural beds of *A. purpuratus* are dispersed along the southwest-American coastline, from northern Peru (Paita) to Tongoy Bay in Coquimbo, northern Chile (1). In Peru, two locations, on the central coast in Independence Bay IB (14.2356 S 76.1929 W) and the northern coast in Sechura Bay SB (5.65 S 80.96 W), are both prominent natural banks.

*Argopecten purpuratus* has become Perús most cultivated marine invertebrate and represents the base of an important aquaculture industry. The capture of *A. purpuratus* spats obtained from natural beds and relocation into fattening areas supports this scallop aquaculture industry. However, in present days, *A. purpuratus* larvae can also be obtained in laboratory under controlled conditions. It is expected that this activity would help to reduce the impact of overfishing of scallop spats in natural beds. However, a critical problem in this intensive larval production of *A. purpuratus* is the recurrent mortality that occurs in controlled systems, often attributed to bacterial infections (2).

Infection outbreaks for *A. purpuratus* larval and postlarval stages are the bottleneck for a successful hatchery of this species. These recurrent bacterial infections cause high economic losses to the marine farming industry, thus, precluding the sustainability of this activity. *Vibrio* species are the primary aetiological agent in *A. purpuratus* pathogenesis disruption (3–5). Reportedly, the vibrios accounted for 98% of the total bacterial isolates from gonads of *A. purpuratus* populations in northern Chile (6).

Scallop larval mortality associated with *Vibrio* and *Pseudomonas* species was described first in 1959 (7), and as a means to control such outbreaks, antibiotics were commonly used. In present-days, probiotic agents are employed as an environmental-friendly alternative to control bacterial infections in aquaculture (8). Therefore, understanding the host-microbe interaction of marine bivalves is paramount for better managing the aquaculture of scallops.

The importance of the interaction between microbes and the corresponding vertebrate and invertebrate host is nowadays well accepted. Hence, some species of bacteria have been found to play a fundamental role in the nutrition and growth of larvae on different cultivated aquatic species, e.g., fish (9, 10) and bivalves (11–14). Beneficial microbiota associated with adult marine organisms are essentially acquired in their early life stages, (15–18). However, the final bacterial species composition may be strongly related to the food style and environmental conditions (19–21).

Among the molluscs, scallops, due to their filter-feeding behavior commonly harbor a complex of associated bacterial community composed of many species within their intestinal tract. Some of them are important as producer of essential vitamins such as B_12_, α-tocopherol, β-carotene, and pantothenic acid, which are indispensable for the correct development and health of scallops larvae (22). Furthermore, it has been demonstrated using isotopic labeled molecules that the scallop *Crassostrea gigaś* larvae can also ingest and digest bacteria, (14, 23). Therefore, the microbial assemblage observed within the adult scallop gut may be strongly associated with the early ingestion of bacteria (24). The complexity of the microbiota associated with bivalves has been largely well-established, diverging significantly in its taxonomic composition (25, 26), which includes both beneficial and deleterious organisms. Reportedly, some Vibrios strains improved the survival of *A. purpuratus* larvae in hatchery systems (11, 27). In contrast, strains of other well know species, e.g., *Vibrio tubiashii*, *V. anguillarum, V. parahaemolyticus,* and *V. alginolyticus* can be devastating for larval cultivation of *Argopecten purpuratus* (3, 5, 28, 29).

The use of commercial probiotics for improving the hatchery of *A. purpuratus* larvae is still questionable. The reason is that the high specificity between microbial-hosts associations could prevent or reduce the successful usage of non-specific commercial probiotic formulations on non-related organisms. As an example, in white shrimp hatchery, the presence of the indigenous bacteria *Halomonas aquamarina* helps to increase the survival of its host *Litopenaeus vannamei,* reducing the pathogenicity of *V. harveyi* (30); however, the mode of action of this interaction is not yet well defined. Therefore, exploring the cultivable microbiota of different marine invertebrates is the first step for isolating species-specific strains. Further, these isolates may be employed as a multispecies formulation of species-specific probiotics, thus improving the results of the low-efficient monostrain probiotics use (31).

To our knowledge, no studies have been conducted to assess the extensive microbial cultivation associated with the scallops in Peru. As part of a survey on the microbiome of the Peruvian scallop *A. purpuratus*, we collected samples from two important growing locations of this species. The main goal of this project is to compare the diversity of culturable bacteria isolated from the intestinal tract of *A. purpuratus* growing on different standard media cultures.

## MATERIAL AND METHODS

### Sampling area and scallop gut dissection

Adult scallops were collected from two locations on the northern and southern coast of Peru: Sechura Bay SB (5.65 S 80.96 W), and Independence Bay IB (14.2356 S 76.1929 W), Fig. 1. A total of 20 adult scallops were captured by autonomous diving at each location at an average depth of 10 m and were maintained in a 20 L tank filled with surface seawater until analysis. The sample size for bacterial culture-dependent enrichment from host invertebrates was determined following the criteria of (32). For this study, ten mature selected scallops were aseptically dissected in the field laboratory, and soft parts (gills and mantle) were carefully removed. Afterwards, the ascending intestine and rectum were traced out as much as possible, then opened scallops were rinsed with particle-free sterile saline solution to clear attached debris, washing out possible contamination from other parts of the organism. Fecal pellets were taken from the distal part of the intestine by flushing out content with 0.5 ml of particle-free sterile saline solution + 20% glycerol (v/v) using a 23-gauge needle attached to a 1 ml syringe, into 5 ml screw-cap tubes.

**Figure 1.**
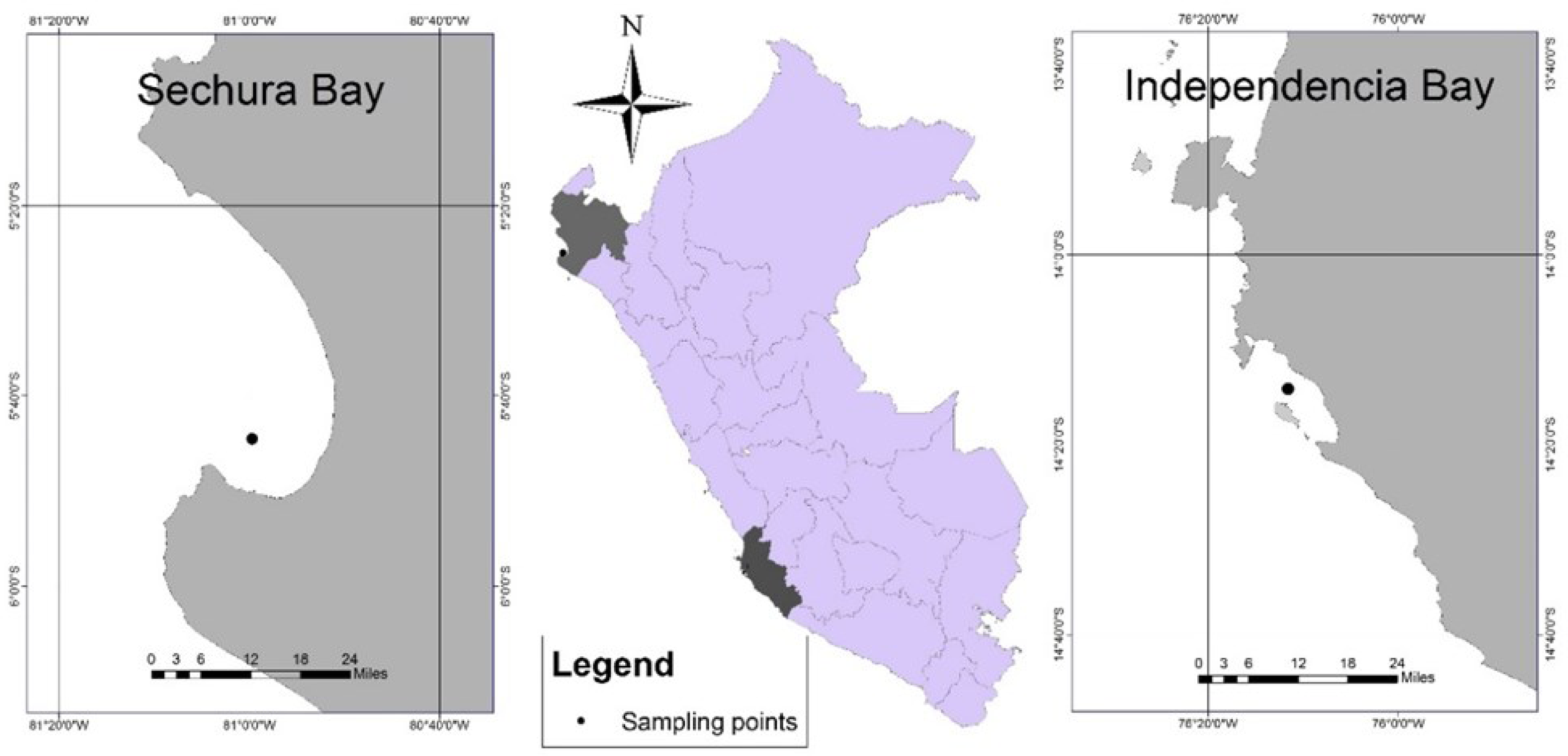
Location of the two *Argopecten purpuratus* main culture areas in the north (Sechura Bay) and central-south (Independence Bay) studied along the Peruvian coastline.

### Enrichment and isolation of cultivable bacteria

For this experiment, 100 µl aliquot (duplicate) of each homogenate was serially diluted (1/10) and plated on: Minimal Medium MM containing per liter (KH_2_PO_4_ (0.5g), CaCl_2_.2H_2_O (0.15g), NH_4_Cl (0.34g), NaCl (30g), MgSO_4_.7H_2_O (0.26g), vitamin B_12_ (1ml), trace elements solution (1ml), agar 15g)); Nutrient Agar (Merck, Darmstadt, Germany); Marine Broth (ZoBell 2216E, Himedia, India), Plate Count Agar, and Czapek Agar, and other selective media such as Man Rogosa-Sharpe MRS, (Merck, Darmstadt, Germany),and Cetrimide agar plates (Merck, Darmstadt, Germany). Samples were incubated at 30°C in the dark for 24 h to 3 weeks. Enrichment growing in MRS medium was incubated in microaerophilic conditions. For isolation of Vibrios, a pre-enrichment using Alkaline Peptone Water (APW) broth was performed; after 12 h incubation, (1/10) diluted samples aliquots were plated on Thiosulfate Citrate Bile Sucrose TCBS agar.

Colonies were selected from agar plate enrichments based on their phenotypic characteristics, color, shape, size, and cell Gram staining. Bacterial pure cultures were obtained by streaking twice a single colony onto fresh agar plates, then one isolated colony was inoculated in a liquid medium to obtain suitable biomass. For long-term storage, active liquid growing cultures were washed twice with sterile particle-free PBS 1X by centrifugation; further, each pure culture pellet was re-suspended with corresponding medium supplemented with 20% (v/v) glycerol (Merck, Darmstadt, Germany) and stored at –80°C.

### DNA extraction and rep-PCR BOX A1R fingerprinting

Bacterial pellets from each isolate were obtained by precipitation at 10000 rpm for 5 min in a centrifuge (Ohaus, USA). Genomic DNA was extracted from the pellets using the Bacterial Genomic DNA Isolation Kit (Norgen Biotech, CA), following the manufactureŕs recommendations. Rep-PCR experiments were performed using the primer BOX A1R (33). PCR reaction mixture containing DreamTaq PCR Master Mix 2X (ThermoFisher Scientific USA), primer BOXA1R (10 µL), and 20 ng genomic DNA (1µl) was completed with PCR grade water to a final volume of 25 ul. PCR reaction conditions were as follows: 5 min at 95°C for initial denaturation, 35 cycles (denaturation 94°C, 1 min; annealing 50°C, 1 min; extension at 65°C, 5 min.). PCR amplified product (5 µl) was used for electrophoresis on a 2% agarose gel in Tris/borate/EDTA (TBE) buffer at 75 V for 120 min, and the gel was stained with SybrGreen fluorochrome mixture (Safe-Green &Trade, ABM Inc. USA). The resulting bands were visualized with a Mini Blue light transilluminator (Cleaver Scientific, UK.) using a 100 bp DNA ladder (Thermo Fisher Scientific, USA) as a molecular weight marker. The DNA banding patterns produced by rep-PCR BOX A1R from isolates of both localities were analyzed by visual examination. All bands were manually scored for the presence (+) or absence (-) to generate a matrix that could be analyzed further. The percent similarity of the banding patterns was estimated with the Dicés coefficient, and the average-linkage-between groups UPGMA (unweighted pair-group method using arithmetic averages) was calculated by the software GelJ (GNU General Public License version 2.0) (34). For instance, single colonies growing on TCBS agar plates with Gram-negative short curved rod shapes were grouped together. Then, presumptive *Vibrio* isolates showing different banding profiles based on rep-PCR analysis were selected for further sequencing. The selection was made visually through a dendrogram constructed with a matrix of presence and absence of amplified bands on agarose gel.

### 16S rRNA gene sequencing and phylogenetic analysis

From extracted genomic DNA of selected strains, PCR was achieved with a reaction mixture containing 2X DreamTaq PCR Master Mix (Thermo Fischer Scientific USA), with 0.5 pmol µl-1 of each primer GM3 (5’-AGAGTTTGATCMTGGC-3’) and GM4 (5’- TACCTTGTTACGACTT-3’) plus genomic DNA template (1 µl) to a final volume of 20 µl, using PCR condition according to (35). Products of PCR amplification (2 µl) were electrophoresed in 1% agarose in 0.5X Tris/acetate/EDTA TAE buffer at 120V for 30 min, and the corresponding size bands were visualized in a Mini Blue Light Transilluminator (Cleaver Scientific UK) using a 1 kb DNA ladder (Thermo Fisher Scientific, USA) as a molecular marker. Prior sequencing the complete ribosomal 16S rRNA gene, a first approach was made by sequencing the short internal fragment of the bacterial 16S with the primer GM1R (5’-ATTACCGCGGCTGCTGG-3’). Then, sequence similarity of each isolate was compared with the GeneBank reference database of sequences using the Basic Local Alignment Sequence Tool BLAST (36). Isolates showing sequence similarity of 100% were grouped together, then one or two isolates from each group were selected for further sequencing of the 16S rRNA gene with the use of additional internal primers: 27F (5’-AGAGTTTGATCMTGGCTCAG-3’) 518F (5’- CCAGCAGCCGCGGTAATACG-3’), and 800R (5’-TACCAGGGTATCTAATCC-3 ’). PCR products were purified and Sanger sequenced at the Macrogen sequencing service (Macrogen Inc. Korea). Resulting chromatograms were analyzed and assembled using the software CHROMAS PRO v2.1.0 (Techlesylum AU). From the chromatogram, the correctness of nucleotide position within the assembly was verified by analyzing the consensus of at least two independent overlapping sequences. The average size of the final assembled sequences was around 1400 bp. GenBank and EZTaxon server was used to identify *de novo* assembled sequences to reference closest relative sequences (36, 37).

16S gene sequences were deposited in the GeneBank database (NCBI accession numbers are listed in tables S2 and S3).

For tree reconstruction, 16S rRNA sequences were also retrieved from the recent actualized and curated database LTP “the all species living tree” project https://imedea.uib-csic.es/mmg/ltp/, based on its recent publication (38). Assembled sequences were first aligned against the fasta formatted compressed LTP database, then the selected closely related reference sequences were realigned using the Multiple Sequence Alignment online service MUSCLE https://www.ebi.ac.uk/Tools/msa/ and further edited and trimmed when necessary, using BioEdit v7.2.5 software (39). The presence of chimeras within the newly assembled sequences was evaluated using the online program DECIPHER web tool http://decipher.cee.wisc.edu/FindChimeras.html.

Phylogenetic trees were reconstructed using the Molecular Evolutionary Analysis (MEGA) package v6.0 (40) through distance-based Neighbor-joining method. Pairwise distances between sequences were evaluated using the Kimura 2-parameter model (41). Maximum-Likelihood (ML), a character-based method for determining phylogenetic inference was also used. ML analysis, including 1000 bootstrap replicates, was performed using RAxML-NG (42) at the CIPRES Science Gateway server version 3.3 (43). Reconstructed trees were visualized with the Interactive Tree of Life (iTOL) server version 5.0 (44).

## RESULTS

### General information about sampling sites

Two natural banks of the scallop *A. purpuratus* located in the northern Peru Sechura Bay SB (5.65 S 80.96 W), and Independence Bay IB (14.2356 S 76.1929 W) in the south were visited twice during the austral spring 2015 and autumn 2016 (Figure 1). SB is located in a transition zone in which the Pacific Equatorial Counter current (warm) collides with the Humboldt current upwelling system (cold). Human activities highly influence SB. Indeed, Parachique, a little fishing dock is located in the southern part of the bay nearby the estuary of Virrilá which is the primary source of freshwater into the bay. Most importantly SB is the main Perús fishing bank where intensive aquaculture production of *A. purpuratus* occurs. Reportedly, SB alone produces more than 50% of the total production of this resource in Peru, according to PRODUCE 2020 (45). In contrast, Independence Bay is a naturally protected area located in the central-southern part of Peru. IB is entirely under the influence of the Humboldt Upwelling System with a relatively low human activity. Biotic and abiotic parameters such as temperature and pH in both locations are shown in the supplementary Table S1.

### Cultivation and phenotypic selection

Bacterial colonies growing in agar plates on different media were selected based on their basic primary characteristics, e.g., form, color, elevation, and margin (Supplementary Figure S1); then grouped accordingly. All tested cultivation media produces different amounts of colonies with different characteristics. The percentage of the relative number of the colonies obtained with each media is shown in Figure 2. As can be seen, the most effective media were the Marine Agar (MA) and Nutrient Agar (NA), which accounted for more than 50% of the total isolated colonies, followed by Agar Plate Count and Minimal Medium Agar with 18.3% and 12.7%, respectively. The other four media altogether produced 18.6% of the observed colonies. TCBS was the most effective selective medium utilized, while Czapek and Cetrimide agar plates were less efficient. On the other hand, no colonies growth was observed in agar plates containing Man-Rogosa-Sharp medium. Additionally, Gram staining was utilized to compare different colony morphotypes.

**Figure 2.**
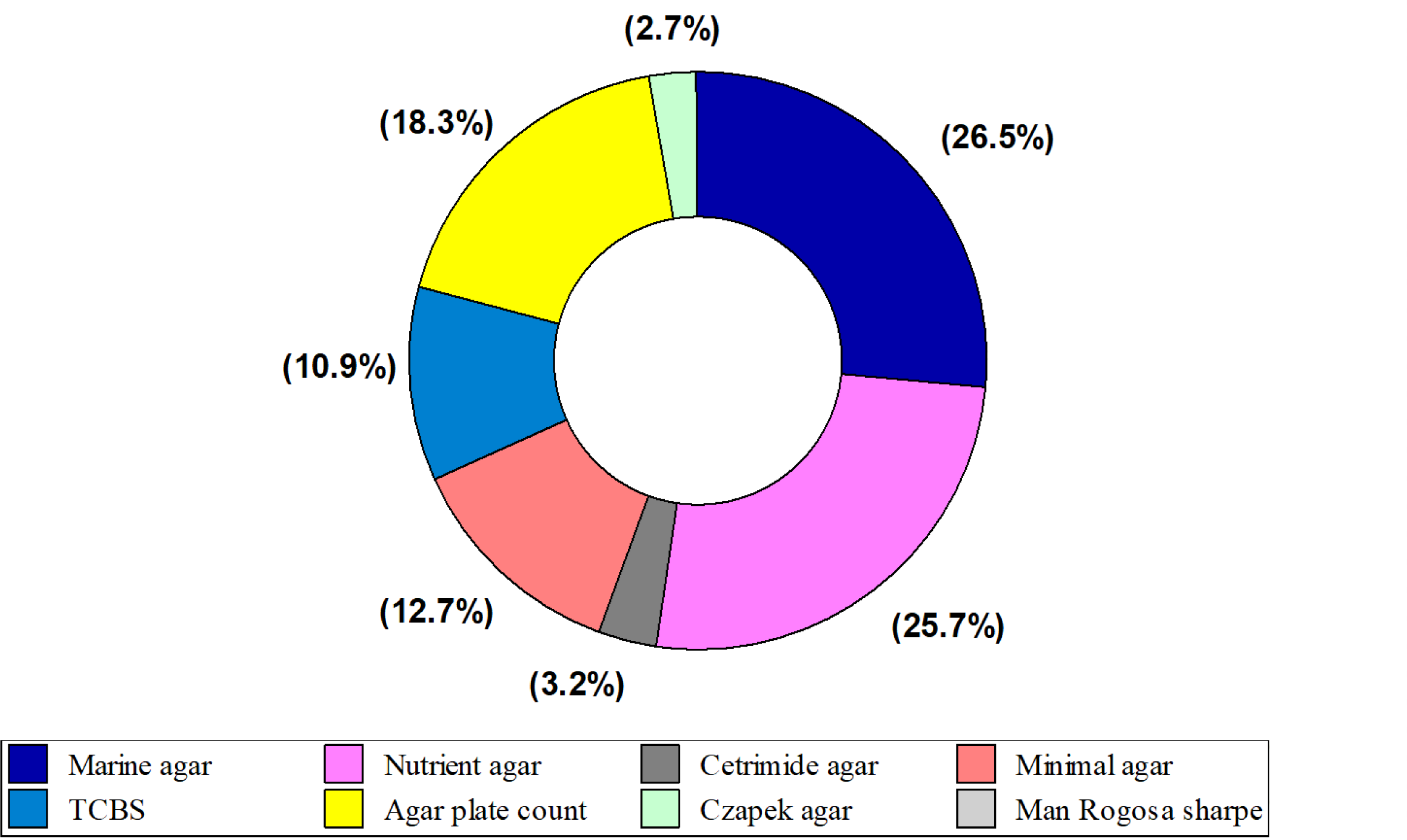
Pay chart showing the relative abundance of strains isolated from the intestinal tract of *A. purpuratus* classified by isolation media (*n* = 339).

### Rep-PCR BOX A1R banding pattern profiles

Following the first phenotypic characterization, purified single colonies obtained by general or selective media were labeled accordingly. Further, isolates that showed similar banding patterns produced by rep-PCR were grouped together. Categorization of Rep-PCR banding profile on agarose gel were made through visual examination. This approach allowed the reduction of the number of isolates for further 16S rRNA gene sequencing. As can be seen in many cases, similar banding patterns were observed in isolates first classified phenotypically as presumptive *Vibrio* and *Bacillus* (Supplementary Figures S2, S3). Therefore, over extensive BOX A1R rep-PCR experiments, a database of banding patterns was created, then bacterial isolates showing different profiles were selected for further sequencing.

### 16S rRNA sequencing and phylogeny

Based on this result, we identified 92 strains at the species level. Species distribution was as follows: 50 strains (IB) and 42 strains (SB). Taxonomically, the major proportions of organisms related to the Firmicutes 46% and Actinobacteria 28.5% were found in (IB), while the proportion for the same phyla in (SB) were 42.8 and 16.6% respectively (Figure 3). Conversely, species affiliated with the Proteobacteria, representing 38.1% of the total species were found in (SB) while this division contributes 26% in (IB) Fig. 3. Remarkably, among the Proteobacteria, the Vibrios originated in (IB) were more diverse than *Vibrio* strains isolated in (SB) represented only by two species. (Table 1 and Supplementary Tables S2 and S3). On the other hand, although the number of strains of the genus *Bacillus* of the Firmicutes were equally distributed in both locations, the species composition differs dramatically being a common representative the species *Bacillus altitudinis*. On the contrary, the actinobacteria were the more abundant and diverse in IB with 14 species contrasting with seven species found in SB (Supplementary Tables S2, S3). Common species in both localities are mostly related to the genera *Bacillus, Priestia, Niallia, Peribacillus*, and *Staphylococcus* of the Firmicutes, whereas the species *Vibrio alginolyticus* was the typical common representative of the Proteobacteria (Figure 4). The topology of the phylogenetic tree constructed with ML, NJ and Bayesian based methods were consistent with each other showing a similar feature (Figures 5A, 5B, 5C).

**Figure 3.**
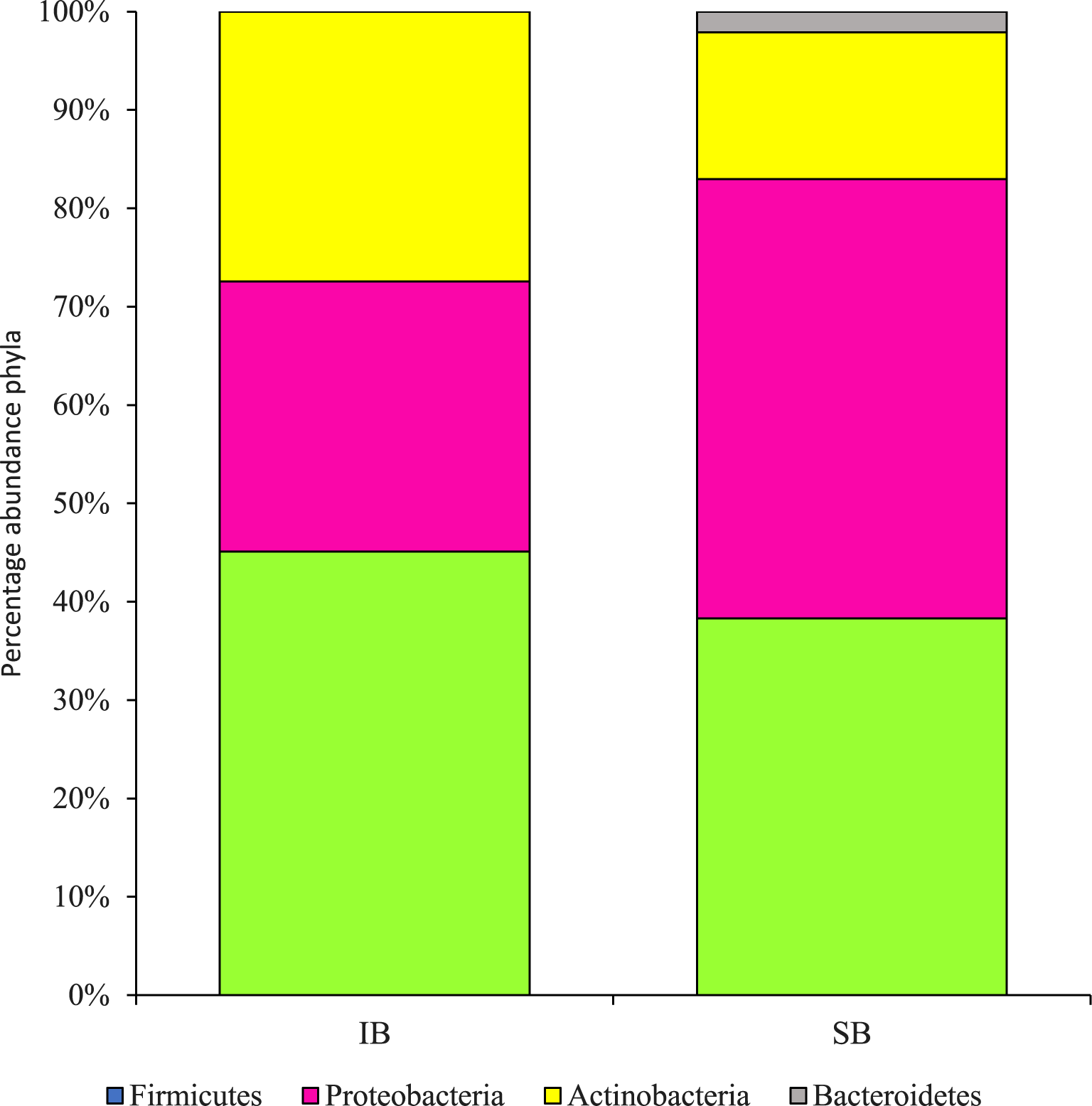
Bar chart showing the percentage of total bacterial isolates identified to species level cultivated in this work. Color indicates the major Phyla, Firmicutes (green), Proteobacteria (red rose), Actinobacteria (yellow), and Bacteroidetes (gray).

**Figure 4.**
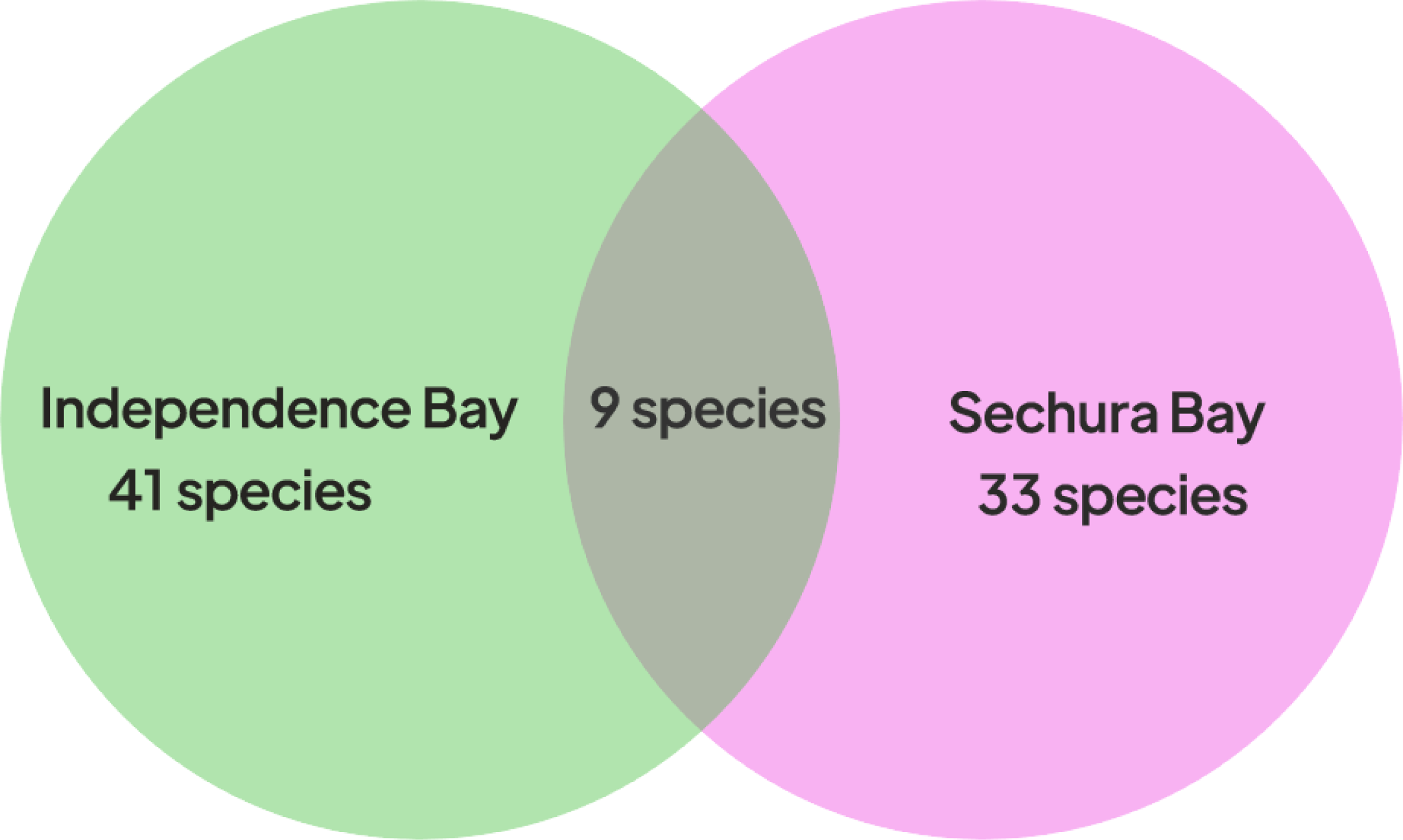
Number of species in common find in two sampling locations in IB (green) and SB (pink red). Venn diagram showing the shared species is represented in grey.

**Figure 5A.**
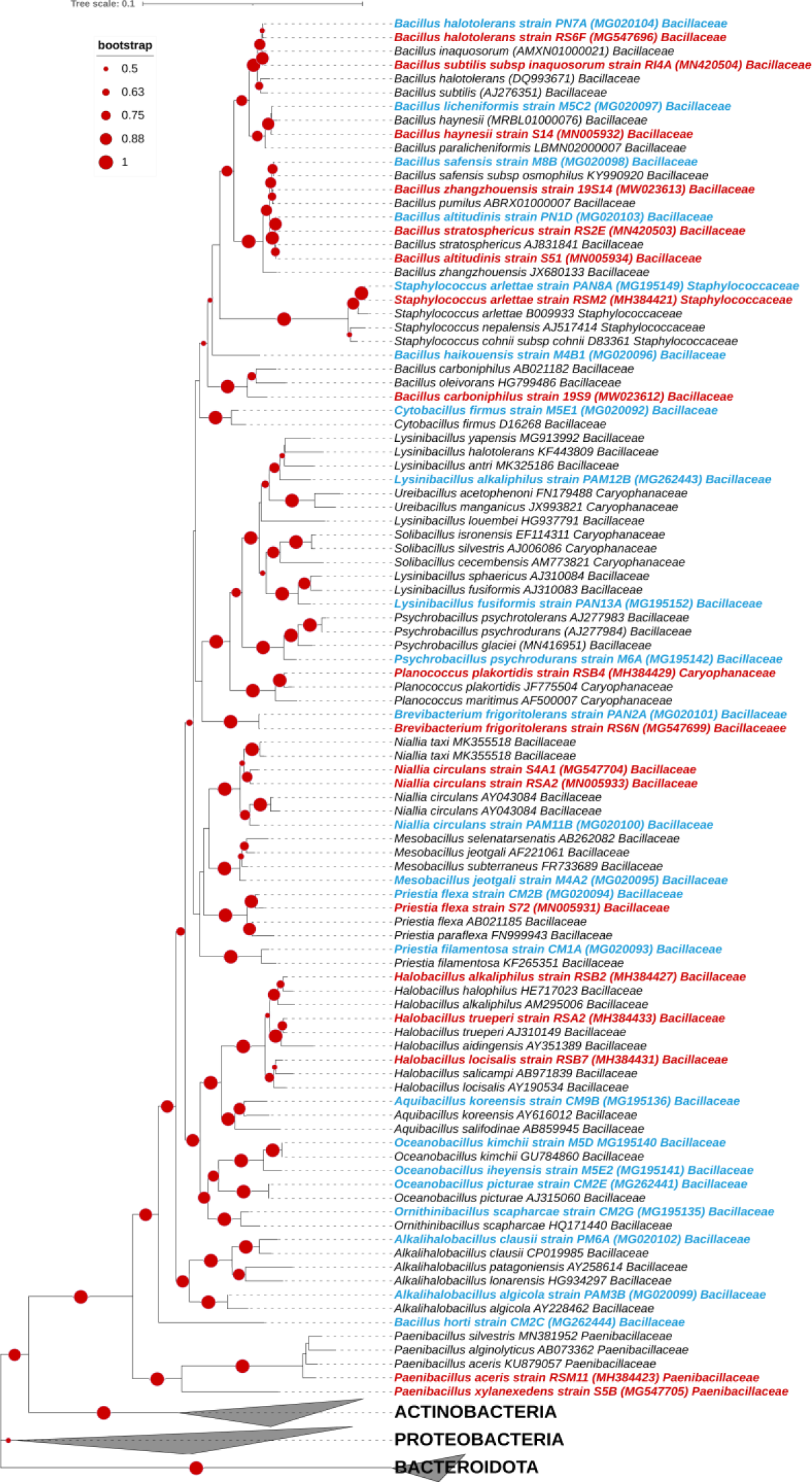
Maximum Likelihood phylogenetic tree of 265 (16S rRNA) nucleotide sequences. Data include BI (blue) and BS (red) sequences of bacteria isolated in this work, close related reference sequences from curated database appears in black. Bootstrap support value >50% is shown (red dots). The scale bar indicates the percentage number of substitution per site. The expanded tree shows the sequences of the phylum Firmicutes.

**Figure 5B.**
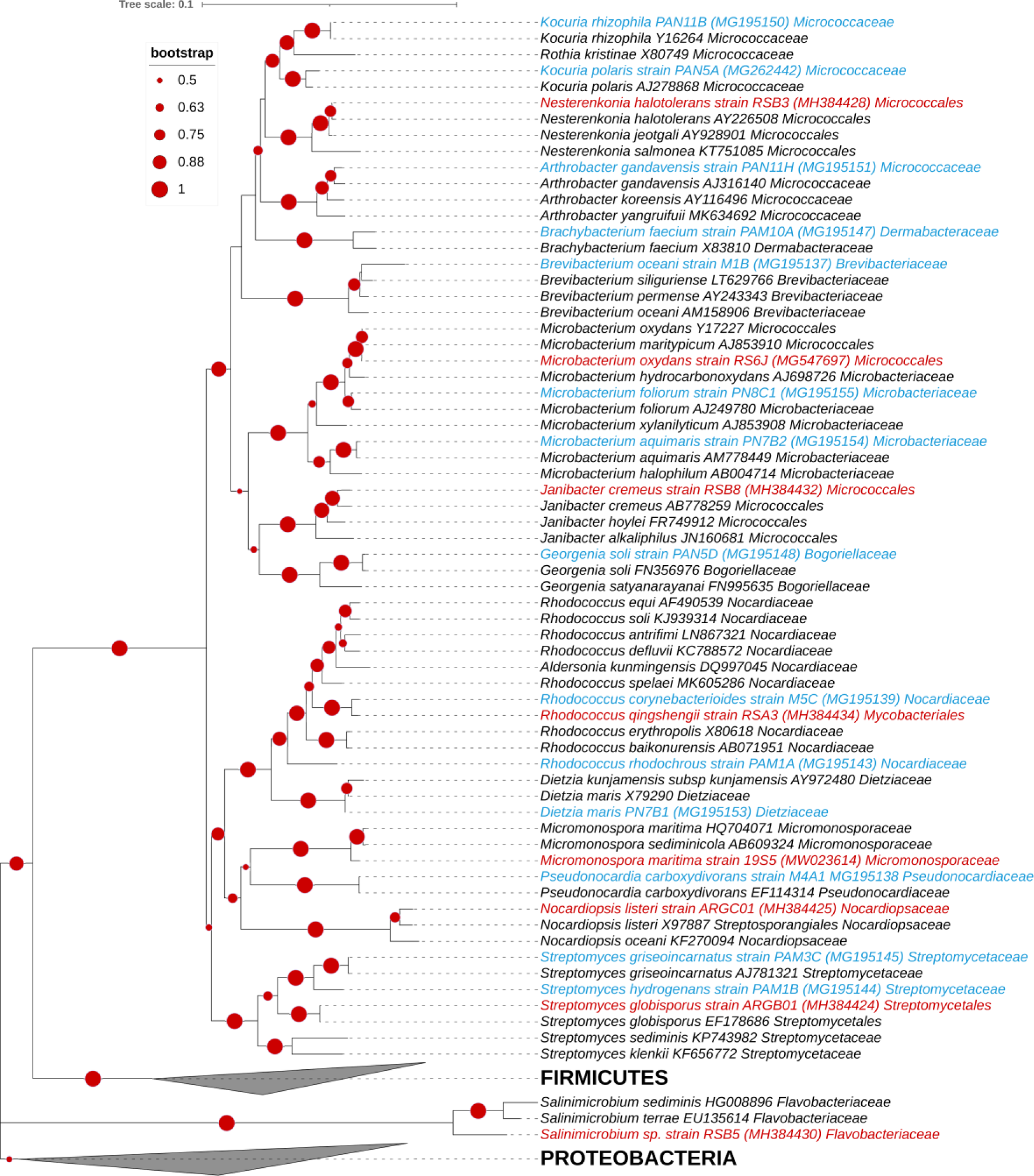
Maximum Likelihood phylogenetic tree of 265 (16S rRNA) nucleotide sequences. Data include BI (blue) and BS (red) sequences of bacteria isolated in this work, close related reference sequences from curated database appears in black. Bootstrap support value >50% is shown (red dots). The scale bar indicates the percentage number of substitution per site. The expanded tree shows the sequences of the phyla Actinobacteria and Bacteroidetes.

**Figure 5C.**
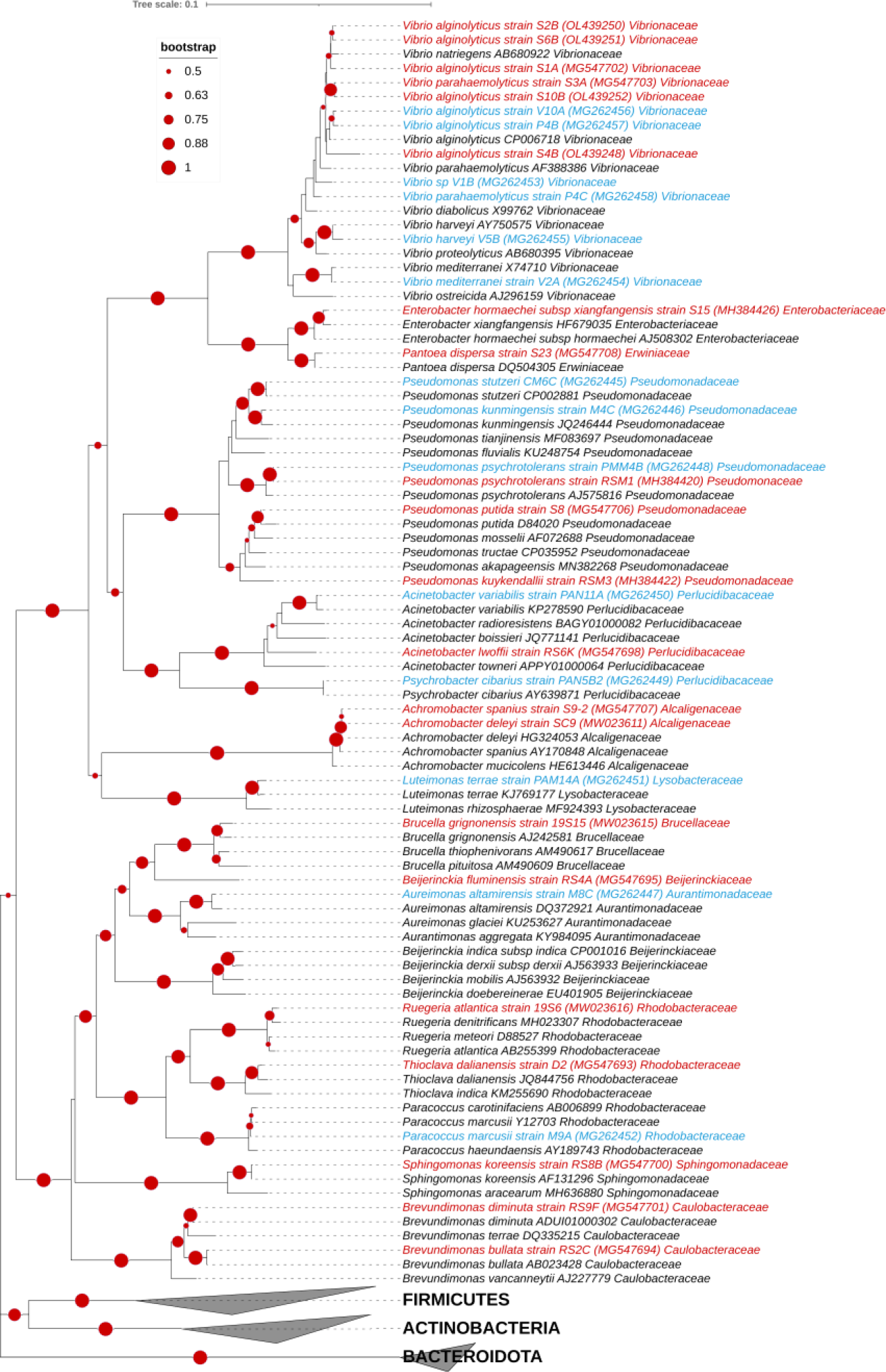
Maximum Likelihood phylogenetic tree of 265 (16S rRNA) nucleotide sequences. Data include BI (blue) and BS (red) sequences of bacteria isolated in this work, close related reference sequences from curated database appears in black. Bootstrap support value >50% is shown (red dots). The scale bar indicates the percentage number of substitution per site. The expanded tree shows the sequences of the phylum Proteobacteria.

**Table 1.** Number of species belonging to each genus is shown.

| Genus | Locations |  |
| --- | --- | --- |
|  | Paracas | Sechura |
|  | (IB) | (SB) |
| *Achromobacter* | 0 | 2 |
| *Acinetobacter* | 1 | 1 |
| *Aquibacillus* | 1 | 0 |
| *Alkalihalobacillus* | 2 | 0 |
| *Beijerinckia* | 0 | 1 |
| *Arthrobacter* | 1 | 0 |
| *Aureimonas* | 1 | 0 |
| *Bacillus* | 7 | 8 |
| *Brachybacterium* | 1 | 0 |
| *Brevibacterium* | 1 | 0 |
| *Brevundimonas* | 0 | 2 |
| *Brucella* | 0 | 1 |
| *Cytobacillus* | 1 | 0 |
| *Dietzia* | 1 | 0 |
| *Enterobacter* | 0 | 1 |
| *Georgenia* | 1 | 0 |
| *Halobacillus* | 0 | 3 |
| *Janibacter* | 0 | 1 |
| *Kocuria* | 2 | 0 |
| *Luteimonas* | 1 | 0 |
| *Lysinibacillus* | 2 | 0 |
| *Mesobacillus* | 1 | 0 |
| *Microbacterium* | 2 | 1 |
| *Micromonospora* | 0 | 1 |
| *Nesterenkonia* | 0 | 1 |
| *Niallia* | 1 | 1 |
| *Nocardiopsis* | 0 | 1 |
| *Oceanobacillus* | 3 | 0 |
| *Ornithinibacillus* | 1 | 0 |
| *Paenibacillus* | 0 | 2 |
| *Pantoea* | 0 | 1 |
| *Paracoccus* | 1 | 0 |
| *Planococcus* | 0 | 1 |
| *Priestia* | 2 | 1 |
| *Pseudomonas* | 3 | 3 |
| *Pseudonocardia* | 1 | 0 |
| *Psychrobacillus* | 1 | 0 |
| *Psychrobacter* | 1 | 0 |
| *Rhodococcus* | 2 | 1 |
| *Ruegeria* | 0 | 1 |
| *Salinimicrobium* | 0 | 1 |
| *Sphingomonas* | 0 | 1 |
| *Staphylococcus* | 1 | 1 |
| *Streptomyces* | 2 | 1 |
| *Thioclava* | 0 | 1 |
| *Vibrio* | 5 | 2 |

## DISCUSSION

The aquaculture of the bay scallop *Argopecten purpuratus* is one of the successful fishery production systems of marine invertebrates in Peru, and is expected to have a stable growth in the following years. Though the extent of populations of this scallop species spread in a patchy way in the occidental south-american coast, SB represents by far one of the most productive areas for the growth of *A. purpuratus* in normal (cold) non “El Niño” conditions. SB alone produces 81% of the total production of *A. purpuratus* in Peru (45). Today, the aquaculture of the bay scallop *A. purpuratus* is the fastest growing food-industry developing in Peru, and it encompasses the world’s growing demand for aquatic food (46). However, at the time this article is written, farm production of *A. purpuratus* in Sechura Bay highly depends on seeds captured in natural beds. Though hatchery production of seeds has experimented a substantial growth, the main drawback of this activity is the recurrent bacterial infections caused primarily by vibrios (3, 4, 5, 47, 48, 49). An alternative to avoid the use of banned antibiotics to control aquaculture diseases is the usage of friendly microorganisms as probiotics, thus requiring species-specific formulation with indigenous organisms isolated from different aquatic cultivated species. In this context, the diversity of the microbiota isolated from the gut of the scallop *A. purpuratus* originated from two natural beds in Peru was studied.

Here, we report the diversity of culturable bacteria obtained through cultivation on general and selective standard media. The scallop *A. purpuratus* is a filter-feeding organism that accumulates a large number of microbes within its digestive tract (24). Although the microbial assemblage can be variable in the different tissues and organs of this organism, the microbiota of the intestinal tract has particularly attracted interest due to the involvement in the nutrition and processing of the food particles captured from the surrounding water.

Currently, the primary approach in studying the bacterial diversity within the intestinal tract of scallops has been the massive sequencing of the 16S rRNA amplicons (50, 51). However, few studies examined the diversity of culturable microbiota. Usually, studies of culturable microflora of *A. purpuratus* have been mainly focused on the identification of pathogenic strains which causes massive mortality in larval hatchery (2, 4, 47, 48). Therefore, this study pinpoints the microflora of *A. purpuratus* targeting the distal portion of the intestinal tract. Our results showed that the cultivated strains predominantly belong to the phyla Firmicutes, Proteobacteria, and Actinobacteria. The Proteobacteria clearly dominate in SB, while Firmicutes moderately predominate in IB. In contrast, the number of species of the phylum Actinobacteria in IB duplicates the number found in SB. Similar phyla distribution was reported by (52) for the oyster (*Crassostrea gasar*), while in the case of culturable bacteria – non-targeting particular organs of the host-isolated in larval cultures of *A. purpuratus* the proportion of each phylum was: Proteobacteria (63%), Firmicutes (24%), Actinobacteria (11%), and Flavobacteria with (2%), (53), in this case, the Proteobacteria appears as a predominant phylum followed by the Firmicutes. In our study, the Firmicutes appears as predominant phyla in IB and SB with 46 and 43% respectively, followed by species of the phylum Proteobacteria and Actinobacteria. However, it is important to remark that the proportion of the Proteobacteria was moderately higher in (SB) than in (IB).

The Virrilá estuary where the artisanal dock of Parachique is placed in the southern part of the Sechura Bay is the main source of freshwater into the bay; therefore, to some extent the bay is under the influence of human activity. Interestingly, two species, *Enterobacter hormaechae* and *Pantoea dispersa* of the family *Enterobacteriacea* were isolated only in SB. The species *P. dispersa* was first reported as commensal bacteria in continental soils, and as a pathogen of plants and insects (54). However, this organism has recently been reported as a potential opportunistic deadly pathogen in humans (55). Conversely, *Enterobacteriaceae-*related strains were absent within the Proteobacteria originated in Independence Bay (IB).

*Pseudomonas putida* strain S8, another suspected opportunistic-related pathogenic species of the Proteobacteria, was also isolated in SB. This metabolically versatile bacterium was isolated together with *P. psychrotolerans* strain RSM1, related to a known cold tolerant species (56), and *P. kuykendalli* strain RSM5 (Table S3). Other *Pseudomonas* species that originated from IB were *P. stutzeri*, *P. kunmingensis* and *P. psychrotolerans* strain PMM4B (Table S2). Despite the involvement of some *Pseudomonas* species, e.g., *Pseudomonas aeruginosa* spp., as an opportunistic human pathogen, pseudomonads are prevalent and are globally distributed in continental and marine environments; hence, due to their versatile metabolism pseudomonads are involved in the mineralization of organic matter, degradation of industrial products including crude-oil, and production of antibacterial metabolites for fish aquaculture (57, 58, 59); therefore, this group of organisms can be helpful in early bioremediation of aquaculture hatchery systems as well as a mean to control *Aeromonas* - a known aquaculture pathogen - proliferation (60). On the other hand, members of the genus *Vibrio* - one of the most conspicuous groups within the Proteobacteria - were also found both in SB and IB. Members of this group of organisms are involved in frequent outbreaks of infections in the aquaculture of the scallop *A. purpuratus* (3, 4, 5, 47). *Vibrio* species are widely spread in aquatic environments, especially in saline habitats. Although pathogenic Vibrios causes enormous losses to the aquaculture industry threatening human health (61, 62), many non-pathogenic species are also recognized to live as symbionts of marine fishes and other invertebrates including the scallops (63, 64). Therefore, benefic *Vibrio* strains can be utilized to control pathogenic strains. Although the mechanism of actions is not fully understood, some benefic *Vibrio* strains possess large cell surface appendages that can prevent the settlement of other related pathogenic strains (65); besides, some strains also showed the potential to produce species-specific antagonist peptides, e.g. bacteriocines (66, 67). Bacteriocin-producing organisms can inhibit other microbial groups growth that share the same ecological niche. Furthermore, in this work IB also showed more diversity of vibrios than SB. (Supplementary Tables S2 and S3).

Regarding the second major phyla, the Firmicutes, the *Bacillaceae* was the most predominant family in both locations in SB and IB. Although the genus *Bacillus* within the *Bacillaceae* is distributed equally in both sites, the number of other genera of the Firmicutes found in IB was 12 compared to 7 found in SB, representing 23 and 18 species, respectively. Here we did not detect the presence of Lactic Acid Bacteria (LAB) related organisms growing in media cultures prepared for this purpose. Reportedly, strains of *Lactobacillus* spp. was frequently recorded as a common probiotic agent in the fish and shrimp aquaculture. (68, 69, 70, 71, 72), but not reported in the aquaculture of scallops. Although non-culturable LAB related organisms were detected in the gut of noble scallop *Chlamys nobilis* (73) using massive sequencing. These amplicon sequences affiliates to the *Lactobacillaceae* of the genus *Streptococcus.* Similarly, following the same method, *Lactobacillus* spp. related sequences were also found within the distal intestinal tract of *A. purpuratus,* thus representing <1% of the total 16S amplicons (50). Conversely, as previously quoted, the genus *Bacillus* within the class *Bacilli* of the Firmicutes was well represented in both locations SB and IB (Tables S2 and S3). *Bacillus* spp. has been extensively reported as a probiotic in fish and shellfish aquaculture (74, 75). The most commonly reported species utilized in aquaculture are: *B. subtilis*, *B. cereus* and *B. pumilus,* all of them were isolated from terrestrial or marine environments (74, 76). However, reviewed literature also listed strains of *Bacillus safensis*, *B. licheniformis*, *B. altitudinis,* and the reclassified species *Nialia circulans* (formerly *Bacillus circulans*), and *Priestia flexa* (formerly *Bacillus flexus*) as probiotic agents (77, 78); strains of these species were also isolated in this study (Supplementary tables S2, S3). Reportedly, strains of *B. licheniformis* and *Priestia flexa* have been isolated from surface sediments of shrimp ponds and other ponds water linked to fish and shellfish farming, and utilized successfully in the aquaculture of white shrimp *Litopenaeus vannamei* (77, 78). Notably, from the above listed isolated strains, the species *Bacillus safensis* is a persistent mesophilic spore-forming bacterium that was first isolated from a spacecraft and assembly-facility system surfaces (79). Here, we isolated a strain of *Bacillus safensis*, the named strain M8B showed a 100% of 16S rRNA sequence similarity with the strain *Bacillus safensis* subsp. *safensis* according to the *EZbioCloud* database (Table S3). *B. safensis*, is a cosmopolitan species found in soil, rhizosphere soil, marine waters, and sediments, as well as in aquaculture water shrimp farms and fish intestinal tracts (80). *B. safensis* is included within the *B. pumilus* phylogroup together with *B. altitudinis* and *B. stratosphericus*. Notably, *B. altitudinis* and *B. stratosphericus* were first isolated in the stratosphere at 41 km above the earth’s surface and reported by (81). In contrast, Lima et al., 2013 (82) reported another *B. stratosphericus* strain LAMA 585 isolated in the Atlantic deep sea at 5,500 m depth. All these species of *Bacillus* are widely distributed on the earth, but instead of *B. safensis, strains of B. altitudinis* and *B. stratosphericus* were found most frequently in marine environments (80). Thus, both species *B. altitudinis* and *B. stratosphericus* here isolated from the gut of the Peruvian scallop *A. purpuratus* could have been originated in marine ecosystems. As was observed in the reconstructed phylogenetic tree, these two species, along with *B. safensis* strain M8B and *B. zhangzhouensis* strain 19S14, clustered together within the *Bacillus pumilus* clade at the top of the reconstructed tree nearby the *B. subtilis* clade Figure 5A.

On the other hand, another bacterial group analyzed in this work was the phylum Actinobacteria. Reportedly Actinobacteria are well-known antibiotic-producer organisms (83). However, although the potential to produce secondary metabolites is widely spread within this phylum, the genus *Streptomyces* has been largely studied as an important source of valuable secondary metabolites. Actinobacteria species found in IB almost duplicate the number of species found in SB (Figure 3). The multi-enzymatic potential and antagonistic compound production by *Streptomyces* can be relevant in aquaculture. Streptomycetes displaying active siderophore production may prevent the establishment of pathogenic vibrios by competing for iron, according to (84). Other modes of action may involve the production of indol-3-acetic acid IAA-like grow-promoting hormone (85, 86) and the production of antimicrobial metabolites (83). In this study, we isolated three *Streptomyces* species from both SB and IB locations (Table 1). *Streptomyces grieseoincarnatus* strain PAM3C isolated in IB, showed a good *in vitro* inhibitory action against the pathogenic *Vibrio harveyii*, *V. alginolyticus* and, the *Bacillaceae Ornithinibacillus scapharcae* (87). Genome analysis of strain PAM3C demonstrates that the organism possesses Lanthipeptide-encoding genes (88); these gene clusters are absent within the genomes of its close related species, *S. variabilis* and *S. griseoincarnatus.* Lanthipetides are ribosomally synthesized derived natural products with antibiotic activity (89).

Bacterial species composition within the gut of *A. purpuratus* from BI and SB varies at different taxonomic levels, e.g., order, family, genus, and species. However, common species found within the intestinal tract of individuals of the two populations were mainly dominated by members of the phyla Firmicutes, with six species shared. In contrast, common species of the Proteobacteria were *Vibrio parahaemolitycus*, *V. alginoltyticus* and *Pseudomonas psychrotolerance.* Important to mention is that none common species were found in the case of the phyla Actinobacteria and Bacteroidota.

On the other hand, considering the distance and discontinuous distribution of *A. pupuratus* natural beds, it was unexpected to find common species between both populations in SB and IB, which are distant by >1000 km each. However, we cannot discard human intervention that illegally translocate scallop spats from distant populations to an environmentally favorable “fattening areas” such as SB (90, 91). Therefore, finding similar “core” species in two disconnected populations of *A. purpuratus* might be a consequence of early bacterial colonization of scallop larvae (24), which still may remain in the adult stage. Moreover, although the number of bacterial cultivated species were almost the same in two populations of *A. purpuratus* collected in SB and IB, the species composition differs significantly. This difference may suggest that the intestinal tract of *A. purpuratus* is a complex community and is indeed strongly influenced by the environment.

As a concluding remark, this is the first study in which extensive cultivation has been made in a localized part of the intestinal tract of the bay scallop *A. purpuratus*. The primary objective of this project was to study the diversity of culturable microbiota of isolates collected in two geographically distant locations. The results obtained in this study may additionally provide primary information for further selection of strains with probiotic capacity. This approach may facilitate the use of species-specific probiotics formulation that ultimately may be helpful to slow the spread of bacterial infection in larval hatchery of the Peruvian scallop *A. purpuratus*.

## Aknowledgements

The authors want to thank the members of the Molecular Microbiology and Bacterial Genomics lab for their valuable collaboration in lab and field assistance. We acknowledge funding from the Peruvian Agency ProInnovate (former InnovatePeru) grant 330-PNICP-BRI-2015 and the Basic Science Research Foundation of the Universidad Cientifica del Sur. Especial thanks to the fisherman association “Asociacion de extractores y maricultores artesanales” of Sechura Bay for allowing us to collect samples in their farming areas, and the “Servicio Nacional de Areas Protegidas” SERNANP of the Ministerio del Ambiente, Peru, for allowing access to the Paracas Natural Reserve, Independence Bay.

## Supplementary material

Supplementary Tables and Figures available on request

## Notes

### Competing Interest Statement

The authors have declared no competing interest.

